# miRSCAPE - Inferring miRNA expression in single-cell clusters

**DOI:** 10.1101/2021.07.29.454389

**Authors:** Gulden Olgun, Vishaka Gopalan, Sridhar Hannenhalli

## Abstract

Micro-RNAs (miRNA) are critical in development, homeostasis, and diseases, including cancer. However, our understanding of miRNA function at cellular resolution is thwarted by the inability of the standard single cell RNA-seq protocols to capture miRNAs. Here we introduce a machine learning tool -- miRSCAPE -- to infer miRNA expression in a sample from its RNA-seq profile. We establish miRSCAPE’s accuracy separately in 10 tissues comprising ~10,000 tumor and normal bulk samples and demonstrate that miRSCAPE accurately infers cell type-specific miRNA activities (predicted vs observed fold-difference correlation ~ 0.81) in two independent datasets where miRNA profiles of specific cell types are available (HEK-GBM, Kidney-Breast-Skin). When trained on human hematopoietic cancers, miRSCAPE can identify active miRNAs in 8 hematopoietic cell lines in mouse with a reasonable accuracy (auROC = 0.67). Finally, we apply miRSCAPE to infer miRNA activities in scRNA clusters in Pancreatic and Lung cancers, as well as in 56 cell types in the Human Cell Landscape (HCL). Across the board, miRSCAPE recapitulates and provides a refined view of known miRNA biology. miRSCAPE is freely available and promises to substantially expand our understanding of gene regulatory networks at cellular resolution.

## Introduction

MiRNAs are small non-coding RNAs that typically bind to 3’ UTR of its target mRNAs and regulate their expression via diverse mechanisms including mRNA degradation^1^, translational inhibition^2^, as well as mRNA stabilization^3^. Additionally, miRNAs can also serve as decoys to indirectly regulate transcription^4^. miRNAs play critical roles in most fundamental cellular processes, from development to homeostasis^2,5^ and consequently are implicated in several diseases, including cancer^6^.

Single-cell RNA sequencing (scRNA-seq) technologies have advanced our understanding of molecular mechanisms at the single-cell level^7^, revealing cellular heterogeneity and identifying previously unknown cellular subpopulations in a variety of contexts^7,8^. However, these technologies are yet to benefit the field of miRNAs, as the reverse transcription stage of the current scRNA-seq protocol relies on poly(A) capture, which mature miRNAs lack. This limitation has precluded the standard scRNA technologies from profiling miRNAs, and consequently, there are only a handful of studies that have profiled miRNA at single-cell resolution^9,10^. The general unavailability of miRNA expression in single cells severely limits our understanding of miRNAs function and dynamics at cellular resolution.

Previous attempts to infer miRNA expression from the expression of protein-coding genes have relied on the assumption that a reduced expression of the putative targets of a miRNA, where the miRNA targets are ascertained based on various sequence-based approaches^11–13^, is indicative of the miRNA activity^14,15^. Such approaches are similar in spirit to SCENIC^16^ which infers activity of a transcription factor in a cell based on the expression of its putative targets. However, due to (i) highly incomplete and noisy nature of miRNA-target relationships, (ii) a highly variable effect of miRNA induction on its targets’ expressions^17^, and as mentioned above, (iii) highly diverse mechanisms underlying the effect of a miRNA on its targets, reliance on putative targets alone to infer miRNA activity is far from ideal; we explicitly demonstrate this assertion. Instead, we hypothesize that the complex direct and indirect regulatory links between miRNAs and the expression of other mRNAs (both protein-coding and non-coding) can be exploited to infer miRNAs at a single-cell level much more effectively than based on putative targets alone. Here, we report a computational tool -- miRSCAPE - to infer miRNAs in single cells from the scRNA-seq data.

First, based on paired miRNA-mRNA profiles in ~5,000 samples in TCGA spanning 10 cancer types and ~4,500 samples in the corresponding healthy tissues in GTEx^18^, we establish the cross-validation accuracy of miRSCAPE in multiple scenarios. Within a test sample, miRSCAPE can accurately rank miRNAs by expression level (average cross-miRNA correlation in predicted and actual ranks within a sample ~ 0.93). For a given miRNA, the correlation between the predicted and the observed expression across the test samples is 0.45 on average, which, as we show in single cell application, is sufficient for an accurate identification of miRNAs that are differentially expressed between two sets of samples. Next, in two independent datasets where cell type-specific miRNA profiles are available (HEK-GBM^9^, Kidney-Breast-Skin^19^), along with the mRNA profiles of those cell types, we show that miRSCAPE, trained on TCGA data, can accurately infer cell type-specific miRNA activities with an average correlation between predicted and observed inter-cell type fold-difference ~ 0.81. Very encouragingly, miRSCAPE is applicable across species; when we trained miRSCAPE on data from 151 samples for human hematological cancer in TCGA and uniformly applied it to 8 hematopoietic cell lines from mice, miRSCAPE significantly distinguished active from inactive miRNAs in each case. Finally, we demonstrate the general utility of miRSCAPE by applying it to scRNA-seq data from Pancreatic Ductal Adenocarcinoma (PDAC), Lung cancers, as well as in 56 cell types in the Human Cell Landscape^7^ (HCL). In each of these applications, miRSCAPE accurately recapitulated the miRNAs previously implicated in each of these contexts and in many cases revealed a cell type-specific role of individual miRNAs.

Overall, miRSCAPE is a versatile and robust computational tool to infer miRNA expression from scRNA-seq data. Application of miRSCAPE to the vast collections of available scRNA-seq will substantially expand our understanding of gene regulatory networks at cellular resolution. miRSCAPE is freely available as a stand-alone tool for the community at https://github.com/hannenhalli-lab/miRSCAPE.

## Results

### Overview of miRSCAPE

Fig 1A illustrates the overall schematic of miRSCAPE. Details are in the Methods section, but briefly, given a large compendium of paired miRNA-mRNA bulk RNA-seq data (Supplementary Table S1), for each miRNA independently, we train an eXtreme Gradient Boosting (XGBoost) model to infer the miRNA expression based on the global mRNA profile of the sample (Methods). We estimate the model accuracy by comparing the predicted and the observed miRNA expression using Spearman rank-order correlation, either within a sample across miRNAs or for a miRNA across samples. In single-cell applications, where both miRNA and mRNA are profiled for multiple cell types, to quantify the model accuracy, we assess the extent to which the cross-cell-type fold-differences of the predicted miRNA expression values are correlated with those of the observed miRNA expression values. Of note, we use all genes as features to train the model instead of using only the experimentally known targets of a miRNA^14^, because, as we show, the model based on all genes consistently outperforms the target-only model (Supplementary results 1).

**Fig 1:**
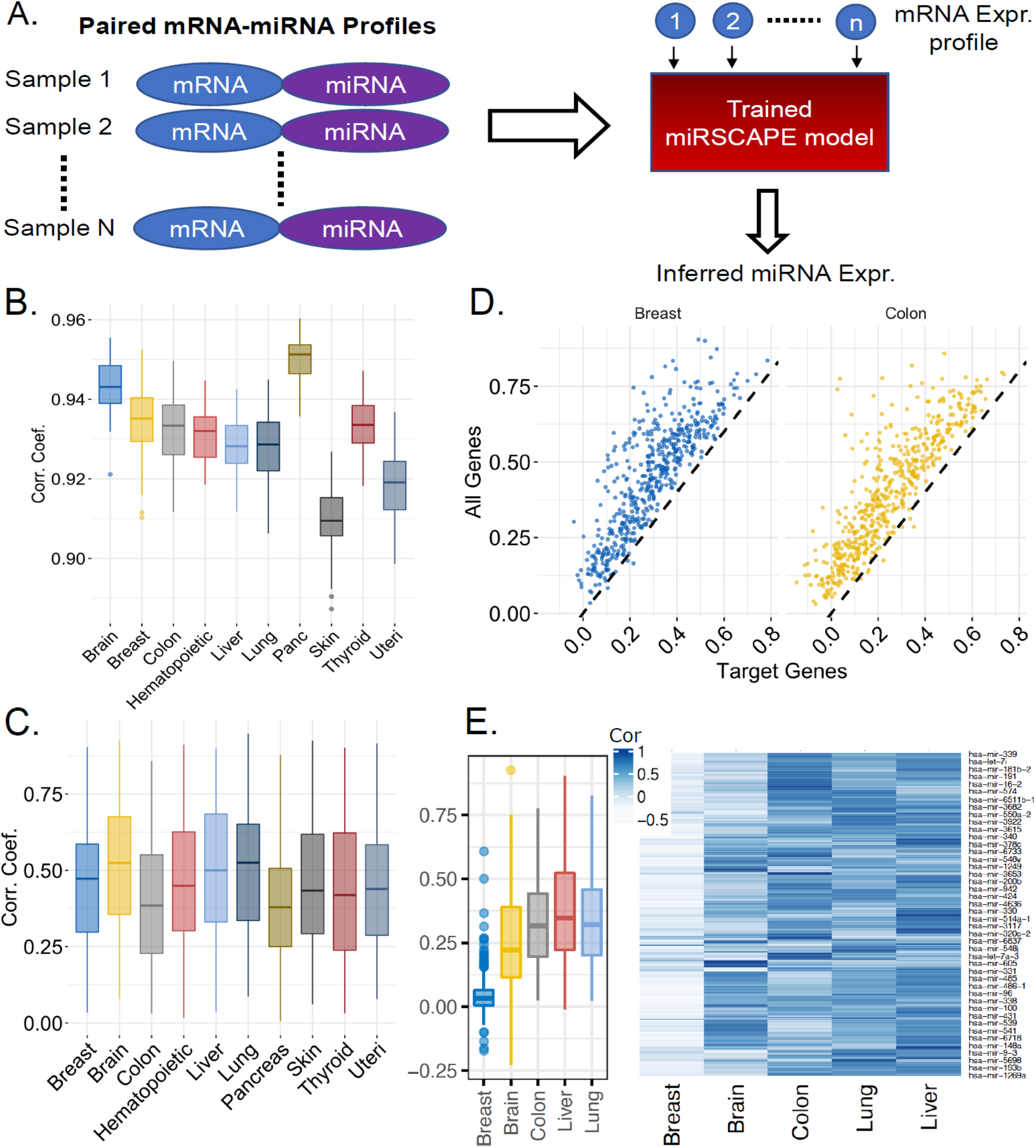
**A. miRSCAPE workflow.** See text for details. B. **miRSCAPE cross-miRNA per sample accuracy in 10 cohorts.** Cross-miRNA spearman rho between the observed and the predicted expressions in each test sample (y-axis) are measured in ten cancer types (x-axis). **C. miRSCAPE cross-sample per miRNA accuracy in 10 cohorts**. Spearman Correlation Coefficients between the observed and the predicted expression across the test samples for each miRNA (y-axis) are calculated in hundreds of samples of ten cancer types in TCGA (x-axis) in a 5-fold cross-validation fashion. **D. miRSCAPE accuracy comparison using all versus target-only genes as features.** In the scatter plots target-only-based accuracy (x-axis) is contrasted with all genes-based accuracy (y-axis) for two tissues; each dot represents a miRNA. **E. Cross-tissue prediction accuracy.** Prediction accuracy when the model is trained on four tissues and applied on the remaining tissue. The heatmap shows the prediction accuracy for each miRNA (rows).

### miRNA Prediction in Bulk Data

We first benchmarked the accuracy of miRSCAPE in cancer bulk paired miRNA-mRNA RNAseq data from TCGA. We selected 10 cancer types from TCGA having at least a hundred matched pairs of miRNA and mRNA data, and in each cancer type independently, we estimated the 5-fold cross-validation (CV) prediction accuracy of miRSCAPE. The miRNAs expressed in fewer than half of the samples within each cancer type were excluded. Within a test sample, miRSCAPE-predicted expression correlates very highly with the observed expressions across miRNA (average Spearman correlation across all tissues ~ 0.93; Fig 1B). When comparing the predicted and observed expression of a given miRNA across test samples, miRSCAPE achieved an accuracy of 0.45 Spearman correlation (Methods) on average, ranging from 0.39 and 0.51 across the 10 cohorts (Fig 1C). Fig 1D illustrates for two cancer types that using all genes as features consistently outperforms the model based on only the known or putative targets, as was done previously^14^; additional information is provided in Supplementary results 1). Additionally, we find a minimal effect of sample size on miRSCAPE accuracy (Supplementary Fig S2).

In many practical instances, there may not be bulk data available to train the model for a specific cell/tissue type in which one wants to predict miRNA expression. To assess miRSCAPE’s applicability in such a scenario, we trained the model using samples from four of the five tissue types and predicted miRNA expression levels in the fifth left-out tissue. Across the five cancer types, and across miRNAs, miRSCAPE achieved an average accuracy of 0.27 (Fig 1E), supporting its broader utility. A relatively lower performance in breast cancer when trained on other cancers is consistent with the fact that the breast cancer samples are transcriptionally the most diverged from other cancers (Supplementary Fig S3).

### Characterizing broadly predictable miRNAs and important gene features

Next, we assessed the extent to which the highly predictable miRNAs and important features (genes) underlying those predictions are shared across cancer tissues. Fig 2A shows the number of highly predictable miRNAs in each tissue at two accuracy thresholds and Fig 2B shows that, of the 10 tissues, on average each miRNA is predictable with rho > 0.3 in 8 tissues and with rho > 0.5 in 5 tissues. miRSCAPE additionally reports the most important features underlying the prediction of each miRNA. First, we assessed whether experimentally known targets of a miRNA are preferentially deemed important by miRSCAPE. Toward this, for a given miRNA, we rank all genes based on the number of tissues in which the gene is identified as an important feature for the miRNA and tested, using Wilcoxon test, whether known targets rank higher than the rest of the genes. Indeed, we found this to be the case for 67% of the miRNAs. Next, we compared, for each pair of cancers, the most important features in each cancer type (those that are deemed important for at least 20% of the miRNAs in the tissue). As shown in Fig 2C, while each cancer has specific important features, there is also a substantive overlap in important features across cancer types (Jaccard similarity ranges from 0.69 to 0.86); only considering the features deemed important for at least 80% of the miRNAs (Supplementary Fig S4), Jaccard similarity remains substantially high ranging from 0.34 to 0.72). Additionally, we observed that tissue-specific important features for a highly predictable miRNA (rho > 0.8) are specifically expressed in the tissue (Supplementary results 2). Lastly, we defined a set of globally important features as those deemed important for at least 20% of the miRNAs in at least 5 cancer types and performed functional enrichment analysis. As shown in Fig 2D, the globally important gene feature genes are largely related to metabolic processes among others.

**Fig 2:**
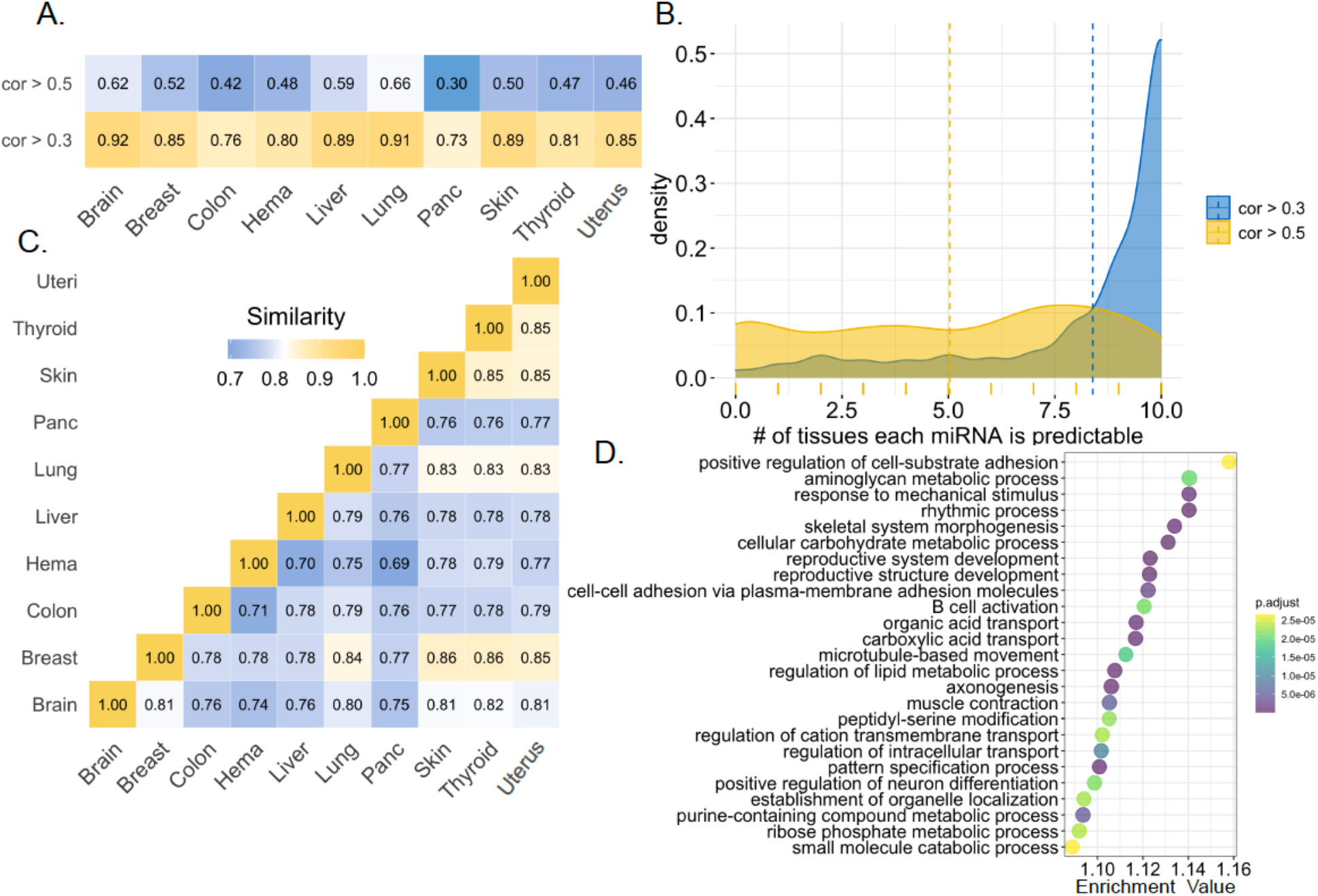
Predictable miRNAs and the contributing gene features. **A.** The fraction of predictable miRNAs for each tissue type at two accuracy correlation cutoffs. **B.** The distribution of the number of tissues (x-axis) in which a miRNA is predictable for two different accuracy correlation cutoffs. **C.** Cross-tissue similarity (Jaccard index) in important features. **D.** Top 25 enriched biological processes among the globally most important gene features.

### Greater sample heterogeneity results in improved model accuracy

To assess the effect of sample heterogeneity on prediction accuracy, we quantified the model performance on 8,089 samples pooled across five cancer types (brain, breast, colon, liver, lung). Fig 3A shows that, across all miRNAs, the pooled model performs substantially better than the cancer type-specific model - the average increase in CV accuracy is 0.2. The improved performance is likely because the model can capture the major differences in miRNA and other gene expression values across tissue types. This is however not explained by the higher sample size in the pooled set (Supplementary Fig S2).

**Fig 3.**
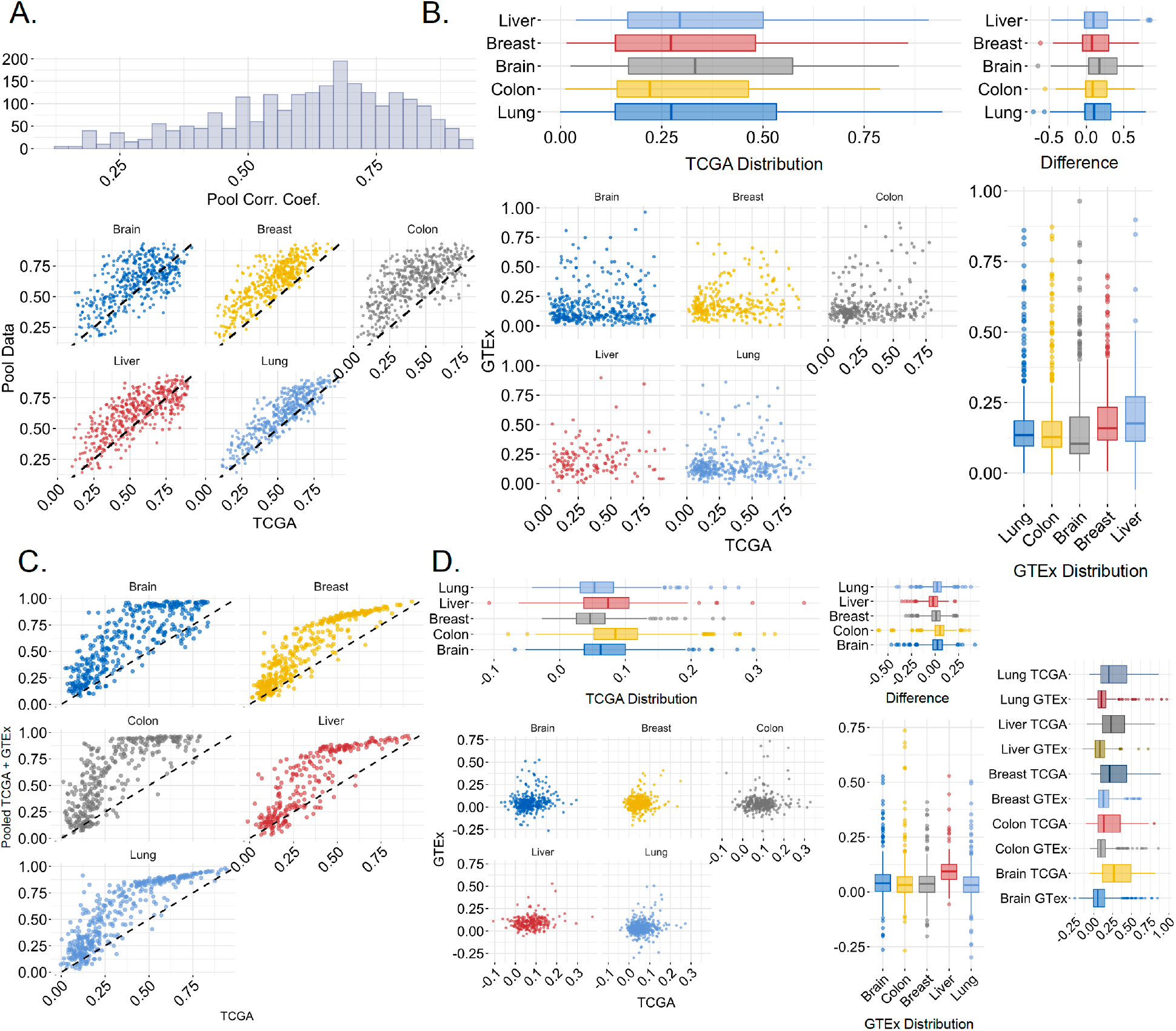
Effects of the inter-sample heterogeneity on model accuracy. **A. Analysis on the pooled samples across five cancer types**. The histogram shows the accuracy for the pooled model, and the scatter plot compares the accuracy for the pooled model (y-axis) with that for tissue-specific models (x-axis); each dot represents a miRNA. **B. Comparison of the cancer tissues with their normal counterparts.** Bottom-left: Scatter plot compares model accuracy for cancer samples (x-axis) to that in healthy samples of the same tissue in GTEx (y-axis). Bottom-right: The boxplots show the distribution of the normal accuracies whereas the boxplots on the top left show the distribution of the cancer accuracies. The difference between the two models is provided in the boxplots on the top right. **C. TCGA versus pooled TCGA + GTEx models.** The scatter plots compare the cancer model accuracy (x-axis) with healthy+cancer pooled accuracy (y-axis). **D. Cross-application of the model.** The scatter plot shows the comparison between models if it is applied to the cross data, the labels represent the data type that the model was trained on. The boxplot on the bottom right shows the distribution of the cross-analysis accuracies when the data is trained on GTEx whereas the boxplot on the top left shows the same when the data is trained on cancer. The difference between the two models is provided in the boxplot on the top right. Rightmost boxplots show the difference between the accuracy when the model is trained and tested on the same data minus the corresponding model tested across data.

Given a greater cross-sample heterogeneity in cancer compared with the healthy counterpart, we compared the prediction accuracies in five cancer types from TCGA with the accuracies in their healthy counterparts from GTEx^18^, as well as with the accuracy on pooled normal and cancer samples. We excluded miRNAs expressed in fewer than 10% of the samples in either the normal or the cancer cohort. As expected, a greater expression variability in cancer results in a more informative model yielding greater CV accuracy (Fig 3B), with an average accuracy of 0.34 in cancer and 0.18 in normal samples. When we pool the normal and the cancer samples, presumably due to differential expression between normal and cancer and further increase in variability, the prediction accuracy is substantially increased to 0.54 on average (Fig 3C); this improved accuracy in the pooled data is consistent with the results obtained above when pooling multiple tissue types (Fig 3A). We further observed that the accuracy of a model trained on cancer samples and tested on corresponding normal samples is substantially better than the converse, again implicating greater heterogeneity in cancer on model accuracy (Fig 3D). However, consistent with our leave-one-tissue-out benchmarking above (Fig 1E), we observed that the cross-cohort accuracy is lower than within-cohort cross-validation accuracy, suggesting substantive transcriptional and regulatory differences across tissues as well as between normal and cancer samples of the same tissue. Overall, these results suggest context-specificity of the model as well as a substantive positive impact of sample heterogeneity on the model accuracy.

### miRSCAPE accurately predicts cell type-specific miRNA expression

Having established the accuracy of miRSCAPE in the bulk data, next we assessed the extent to which miRSCAPE, trained on bulk data, can infer miRNA expression in particular cell types (Fig 4A and Methods). We tested this in three independent datasets where miRNA and mRNA profiles are available for individual cell types based on either single cell profiling or bulk profiling of purified cell types.

**Fig 4:**
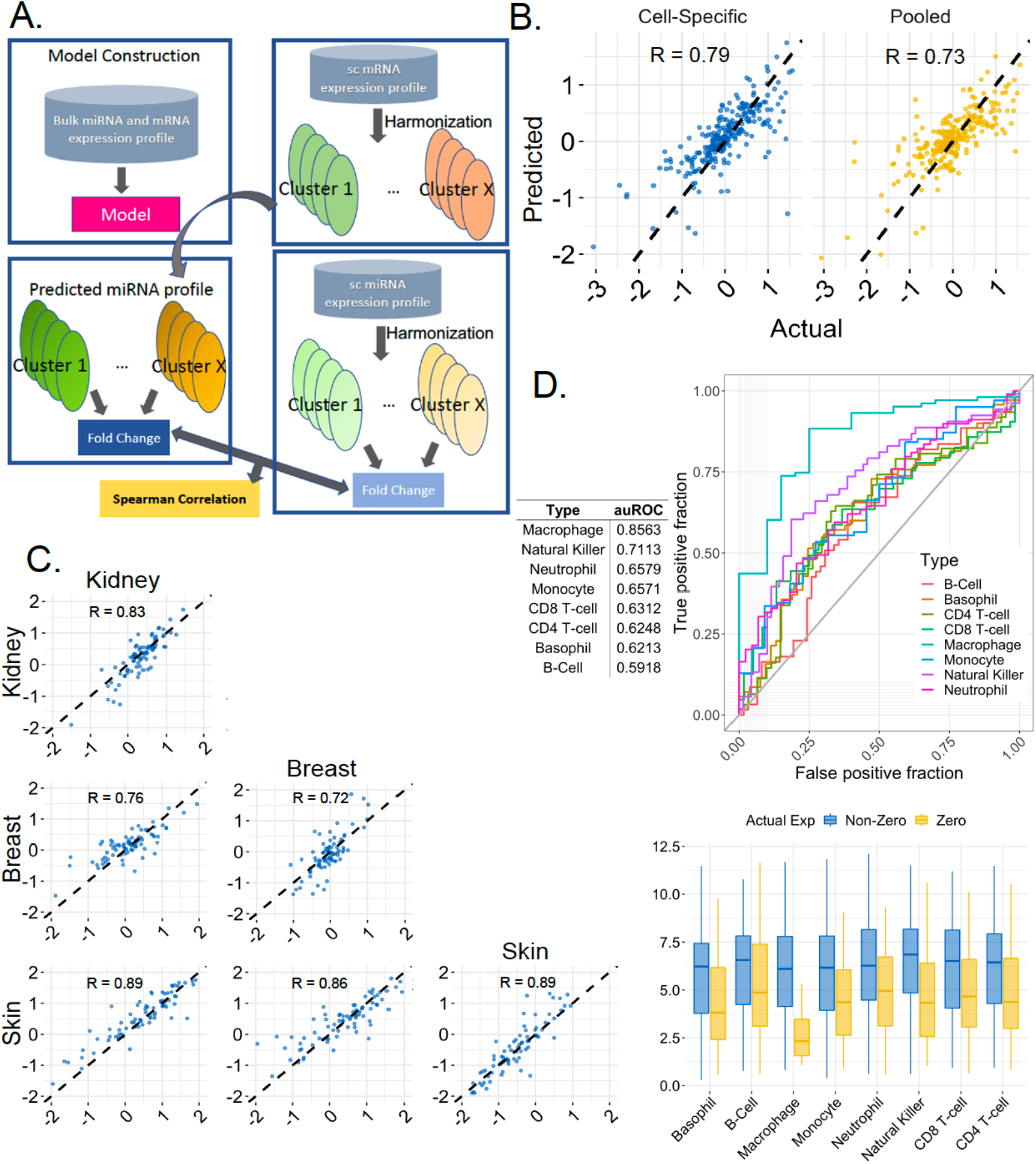
miRSCAPE validation in independent single cell data. **A. Validation pipeline.** A model is learned for the matching bulk data and then miRNA expression is inferred from the single-cell mRNA data for individual cell types. To assess miRSCAPE predictability, for each miRNA, the fold-difference between two cell types using the observed and the predicted miRNA expression are estimated, and the two sets of fold differences are compared across miRNAs using Spearman correlation. **B. Validation on Faridani et al data.** Scatter plots comparing fold-difference across cell types using observed miRNA expression (X-axis) and predicted expression (y-axis). Left plot is based on cell type-specific models, and the right plot is based on a single pooled model. **C. Validation on Isiakova et al.** Refer to B for details. Off-diagonal plots compare two cell types, and the diagonal plots compare one cell type (row) with the other two cell types pooled. **D. Validation on hematopoietic cell types.** Box plot for miRSCAPE prediction values and the corresponding ROC curves and the table for auROC values show the extent to which miRSCAPE could distinguish the miRNAs having experimentally detectable levels of expression from those with no detection in the cell type in 8 major hematopoietic cell types.

Faridani et al.^9^ (GSE81287) measured single-cell miRNA expression profiles in the brain (166 cells) and kidney (45 cells). We trained miRSCAPE separately on the bulk brain and kidney cancers from TCGA and applied the models respectively to the scRNA-seq profiles of brain and kidney cells obtained from the HCL^7^, yielding inferred cell type-specific miRNA profiles. We then compared the kidney vs brain fold-change (FC) based on the predicted miRNA expression with those based on observed miRNA expression from Faridani et al.^9^ Fig 4B shows that across 262 miRNAs, the predicted and the observed FC are highly correlated (Spearman rho = 0.8). Even when we used a single model trained on the pooled brain and kidney bulk data and applied the same model to predict the miRNA expression for both kidney and brain scRNA data, miRSCAPE achieved a high concordance between the predicted and observed FC (Spearman rho = 0.73; Fig 4B).

Isakova et al.^19^ have applied Smart-seq-total simultaneously to profile within the same cell both miRNAs as well as protein-coding mRNAs in the skin (277 cells), breast (90 cells), and kidney (245 cells). We trained cell-type-specific miRSCAPE models for skin, breast, and kidney from the samples for the corresponding cancer types in TCGA (Methods) and applied those models to the scRNA-seq data to predict cell-type specific miRNA profiles in the three cell types. For each cell type pair, as above, we compared the FC between the predicted and observed cell type-specific miRNAs. As shown in Fig. 4C, the Spearman rank correlation coefficient for each of the three comparisons range from 0.76 to 0.89.

While in the above examples we compare highly diverged cell types, next we assessed miRSCAPE’s ability to infer different miRNA expression amongst relatively similar cell types within the hematopoietic system. Since we could not obtain both miRNA and mRNA profiles in human hematopoietic cell types, we instead assessed the extent to which miRSCAPE trained on human bulk data can infer miRNA activities in mouse hematopoietic cell types. Toward this, we trained a miRSCAPE on 151 Acute myeloid leukemia samples in TCGA. In Supplementary results 3, we provide substantiation of our model based on a miRNA knock-out data.

We obtained transcriptional profiles of >15,000 mouse cells across 8 major immune cell types from Zilionis et al. study ^20^ (GSE127465), pooled and harmonized the data for each cell type, and applied the miRSCAPE model to infer cell type-specific miRNA activities. Using purified bulk miRNA-seq data for the 8 hematopoietic cell types^21^, for each cell type individually, we assessed whether the inferred expression of miRNAs could distinguish the miRNAs having experimentally detectable levels of expression in the cell type from the undetected miRNAs. However, since only a relatively small fraction of miRNAs are detectable in a cell type, instead of quantifying accuracy using correlations, we used Wilcoxon test as well as quantified the classification accuracy using auROC. In all 8 cell types, this was indeed the case (Wilcoxon P <= 0.05, average auROC = 0.67) and Fig 4E shows the ROC curves and the auROC values for each cell type, clearly suggesting that a model trained in human bulk data can still identify cell type miRNAs in the mouse.

Overall, in all independent validations in single-cell or bulk purified cell type datasets, our model trained on human bulk data, including cross-species application, achieved a high concordance between the predicted and the observed miRNA expression, firmly establishing the generalizability and robustness of the model.

### miRSCAPE reveals miRNA activities in the tumor microenvironment and across normal fetal and adult cells

Having established the accuracy of miRSCAPE via cross-validation and in multiple independent single cell and cell type-specific datasets, we set out to demonstrate miRSCAPE’s utility in multiple contexts.

### Pancreatic Ductal Adenocarcinoma (PDAC)

We applied miRSCAPE, trained on TCGA PDAC cohort, to scRNA-seq data in PDAC^22^ comprising 57,730 cells from 35 donors. The cell annotations were obtained from Peng et al.^22^ and here, we focused on three cell types - acinar, normal ductal (Ductal type I), and the malignant (Ductal type II) cells. We applied miRSCAPE to each cell type separately on 150 pools of randomly selected 22,225 cells to obtain the pseudo-bulk transcriptome to predict miRNA expression distribution (Methods).

The cell type that gives rise to PDAC is not entirely resolved and both acinar, as well as ductal cells, represent likely candidates for the cell of origin for PDAC^23^. We, therefore, compared the differential miRNA expression in the malignant cells relative to acinar and ductal (Type I) cells individually as well as relative to pooled acinar and ductal cells. Supplementary Table S2 shows the significantly differential miRNAs in the malignant cells in all three comparisons.

Based on bulk transcriptomes, Mazza et al.^24^ have reported differentially expressed miRNAs in PDAC relative to the normal pancreas, of which 39 are included in our study. As shown in Fig 5A and Supplementary Fig S5, our findings based on single-cell data are highly consistent with Mazza et al. -- overall, 24 of the 39 miRNAs show differential expression in the malignant cells relative to either acinar or ductal cells, and the correlation between our predicted and observed fold changes from Mazza et al. is 0.58 on average across all pairwise comparisons. As an example, miR-221, identified by miRSCAPE, is known to be upregulated in pancreatic cancer cell lines^25–27^ and plays a role in invasion, drug resistance, and apoptosis in PDAC^26^. Likewise, overexpression of miR-29a sensitized chemotherapeutic resistant pancreatic cancer cells to gemcitabine, reduced cancer cell viability, and increased cytotoxicity^28,29^. Interestingly, miR-29a has also been shown to have a tumor-suppressive role in PDAC^30^, and based on miRSCAPE, miR-29a is predicted to be downregulated in malignant cells relative to acinar cells but surprisingly, is upregulated relative to ductal type I cells. This suggests a pleiotropic and potentially context-specific role, and also suggesting acinar as the cell of origin for PDAC^31^.

**Fig 5:**
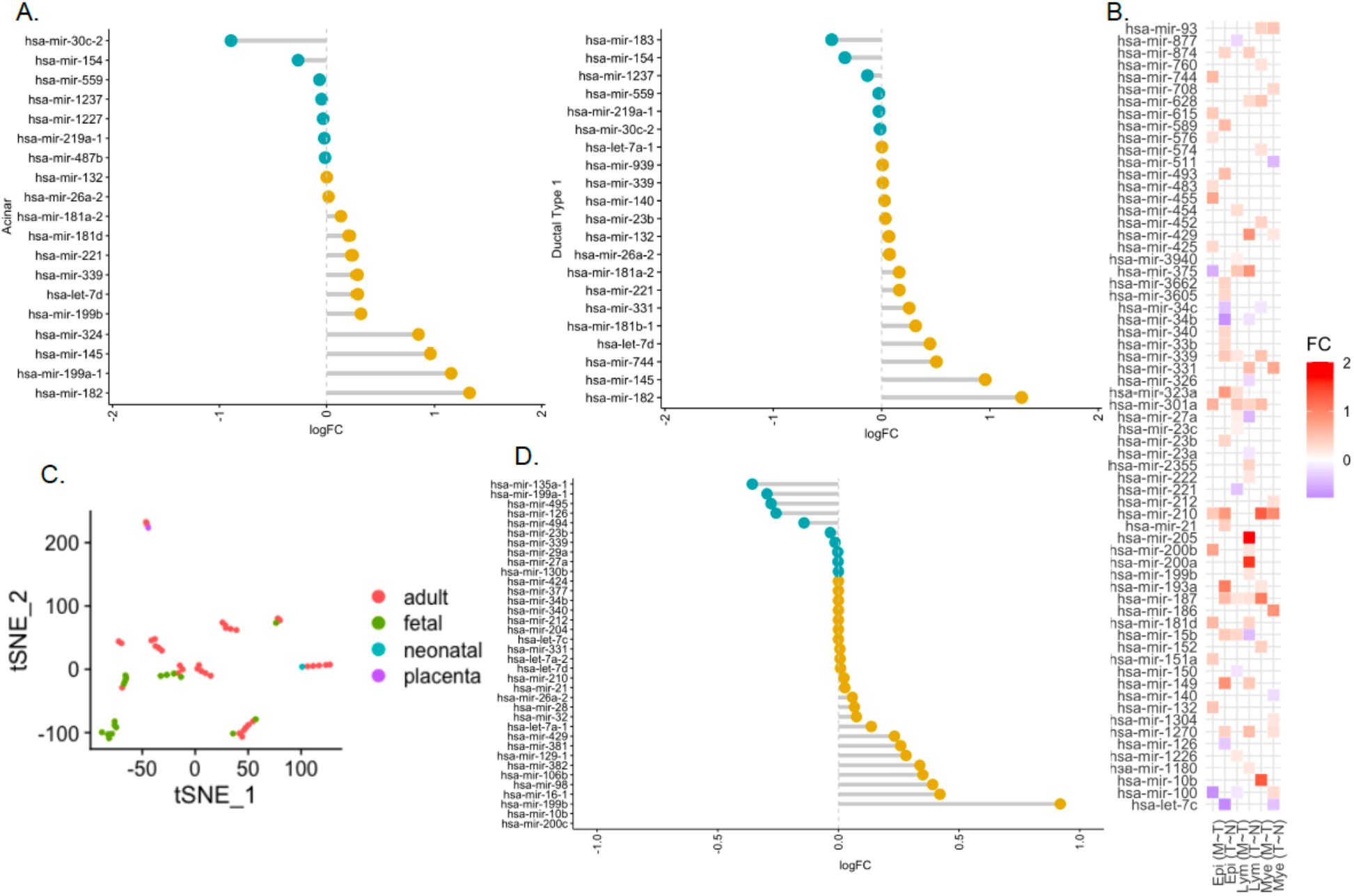
miRSCAPE application to single cell data. **A. Application to PDAC.** Fold change (x-axis) the predicted miRNAs (y-axis) differentially expressed in malignant ductal type 2 relative to normal acinar (left) and ductal (right) cells; miRNAs whose predicted fold-change direction is consistent with Mazza et al. are shown. **B. Application to lung adenocarcinoma.** Heatmap for predicted fold changes of the miRNAs (y-axis) in each of the epithelial (Epi), lymphoid (Lym), and myeloid (Mye) cell types in Primary tumor~Normal (T~N) and Metastasis~Primary tumor (M~T) (x-axis). **C. Application to human cell atlas.** tSNE plot for cell types based on their global predicted miRNA profiles. **D. Fold change distribution (x-axis) of the predicted miRNAs (y-axis) in fetal~adult heart.** miRNAs whose predicted fold-difference is consistent with Thum et al. are shown.

### Lung Cancer

Next, we applied miRSCAPE to scRNA-seq of lung adenocarcinoma^32^ consisting of 208,506 cells derived from 44 individuals including 11 biopsies from adjacent-normal tissues, 14 primary tumors, and 9 metastatic tumors^32^. Cell cluster annotations were obtained from the original publication, and here we focus on three cell types - epithelial, lymphoid, and myeloid cells. For each cell type, separately in normal, primary tumor, and metastatic tumor samples, as above, we pooled randomly selected cells to predict miRNA expression distributions in each cell type.

We compared the primary tumors with normal samples and also the metastatic tumors with the primary tumors separately in epithelial, lymphoid, and myeloid cells and identified the top 20 upregulated and downregulated miRNAs among 402 miRNAs (Supplementary Table S3) for each cell type in each comparison. The union of the top 20 upregulated and downregulated miRNAs in each of the three cell types in Primary tumor~Normal and Metastasis~Primary tumor includes 149 unique miRNAs, 86 of which are listed in the OncomiR database^33^. Of these 86 miRNAs, miRSCAPE identifies 65 to be consistently differentially active in Lung Adenocarcinoma (LUAD) and Lung Squamous Cell Carcinoma (LUSC) cancers, suggesting a high concordance (75.6%) between miRSCAPE prediction and OncomiR (Fig 5B). Broadly, miRSCAPE not only recapitulates many of the miRNAs associated with lung cancer, it in fact reveals their cell type-specific roles (e.g., miR-4664 is upregulated specifically in epithelial cells of the tumor), which in some cases is even opposing in different cell types (e.g., miR-328 is downregulated in primary tumor myeloid cells but upregulated in the brain metastatic lymphoid cells). Many of our detected tumor-associated miRNAs in immune cells are known to be associated with leukemias, e.g., miR-21. We have provided further details of a few interesting cases in Supplementary results 4.

Overall, our results suggest that miRSCAPE not only identifies key miRNAs involved in lung cancer it also provides an opportunity to investigate cell type-specific roles of these miRNAs in the context of lung cancer.

### Global Model

Finally, we set out to chart a comprehensive global landscape of miRNA expression across all human cell types profiled via scRNA-seq in the HCL^7^. Our goal was to train a single global model that captures the co-variation among miRNAs and mRNAs and is uniformly applicable to all cell types. Toward this, we collected tumor and normal samples across ten diverse tissues (Supplementary Table S1), comprising 13,764 samples. To reduce the sample space without compromising the captured variance, for each tissue separately, we clustered the tumor and the normal samples using k-means clustering into 100 clusters. We then selected the medoid of each cluster as its representative, yielding 100 samples for each tissue, with a total of 1,000 samples representing the global variation in the human body. These 1,000 samples were used to build a single global miRSCAPE model.

We obtain scRNA-seq data from HCL^7^, encompassing >700,000 cells across 56 different cell types comprising 36 adult, 18 fetal, 1 neonatal, and 1 placental sample. We pooled and the scRNA-seq profiles for each cluster and applied the global miRSCAPE model to each cell type yielding inferred activities of 523 miRNAs across the 56 cell types. Top 20 upregulated and downregulated miRNAs in each cell type relative to all other cell types are provided in Supplementary Table S4.

We first demonstrated that clustering of cell types based on their global predicted miRNA profiles clearly segregates the fetal from the adult tissues (Fig 5C) and is highly concordant with a similar clustering based on the global mRNA profiles of the 56 cell types, with an overall Spearman correlation of 0.86 across all pairwise cell type distances across the two modalities. Supplementary Table S5 lists the top 20 differential miRNAs between the fetal and the adult tissues of each type. These results are meant as a public resource for a variety of downstream analyses. As such, knowledge of developmentally regulated miRNAs in individual tissues is relatively limited for us to corroborate our findings. Nevertheless, in the few tissues where such data are available miRSCAPE recapitulates previous findings to a reasonable degree (Fig 5D), and is further discussed in the Supplementary results 5.

Overall, in a variety of contexts, miRSCAPE recapitulates the known biology and its application to the 56 cell types in the human cell atlas will serve as a valuable resource for the community to explore the developmental roles of miRNAs in various tissues.

## Discussion

Single-cell RNA-seq technologies have matured and are routinely used to generate large scRNA-seq data, effectively capturing protein-coding genes, in a wide variety of contexts. However, analogous technology specifically for small non-coding RNA sequencing, specifically miRNAs, is significantly lagging and has only been demonstrated in a handful of cell types^9,19^. This has left a substantial gap in our understanding of miRNA transcriptional dynamics at cellular resolution. To overcome this limitation, here we report a machine learning tool, miRSCAPE, to predict the miRNA expression in single-cell clusters from their genome-wide mRNA profiles. We extensively demonstrate miRSCAPE’s accuracy both in a cross-validation fashion in several tumor and normal bulk data sets, as well as in multiple independent single cell data sets. Finally, we show that miRSCAPE can successfully recapitulate differentially expressed miRNAs in Pancreatic Ductal Adenocarcinoma and Lung cancer, as well as in 56 normal cell types in the HCL. Our tool and the associated freely available software open up the possibility to leverage a vast compendium of scRNA-seq datasets to understand miRNA activities at the cellular resolution.

The mechanistic premise of miRSCAPE is that the miRNA activity is reflected, directly or indirectly, in the global transcriptomic profiles. However, the global transcriptomic profile reflects not only miRNA activity but several other cellular features, such as protein activities, metabolite levels, enhancer activity, DNA methylation, etc. and in principle, machine learning can predict these other cellular features from the global transcriptomic profiles. Thus, the miRSCAPE framework can be extended to estimate these other features at a cellular resolution, if appropriate paired data are available for training the model. One limitation of miRSCAPE is that because it is trained on bulk data, it is not expected to perform well if a large fraction of features has missing values, as is the case for single cell transcriptomic profiles. Hence, we pool cells within a single cell cluster and apply miRSCAPE to the pooled (and therefore not sparse) transcriptomic profile to infer miRNA activity at the level of single cell clusters. However, this is not a major limitation because miRSCAPE can still identify miRNA activity for distinct cell types or cell states as long as those types or states are discernable based on the scRNA-seq profile by the standard tool^34^. To add, the notion of a ‘cluster’ is relative, and in a typical scRNA-seq application, one can either define clusters at a very high resolution or even pool nearest neighbors to estimate smoothed miRNA expression values.

miRSCAPE is expected to perform the best when it is trained on a large training set that captures the transcriptional diversity of the specific context it is applied to. However, when a precise cell or tissue type data is not available, one can consider a closely matching context for training. In a situation where multiple cells or tissue types are involved, a global model spanning all such contexts can be useful (as in our HCL application), with a small loss in performance compared to context-specific models, as demonstrated in our HEK-GBM validation (Fig 4B). When the training context is very different from the application context, a more substantive loss in performance is expected, as quantified in Fig 1E. However, when using a global model, loss in performance may be compensated by the ability to uniformly apply a single model, making the inferences in specific sub-contexts directly comparable (Fig 5C, 5D).

Overall, miRSCAPE enables studying the miRNA transcriptional dynamics in a vast variety of contexts where scRNA-seq data has been profiled, from development to homeostasis to diseases including cancer. We have comprehensively benchmarked miRSCAPE and demonstrated its utility in multiple contexts and made the tool freely available. miRSCAPE thus represents an impactful advance toward leveraging the scRNA-seq data to expand our understanding of transcriptional dynamics at cellular resolution. miRSCAPE is available at https://github.com/hannenhalli-lab/miRSCAPE.

## Methods

### Data

We obtained bulk matched RNA-Seq and miRNA-Seq data for cell lines (Cancer Cell Line Encyclopedia, CCLE^35^), normal tissues (GTEx^18^), and cancer (TCGA). We selected ten tissues having more than a hundred matched mRNA and miRNA samples in TCGA and GTex. Experimentally known miRNA target information was gathered from the miRTarBase^36^.

We make use of various single cell mRNASeq and miRNAseq expression profiles. Small seq single cell miRNA expressions are obtained from Faridani et al. (GSE81287) and Isakova et al. (GSE151334) studies. Mouse hematopoietic single cell mRNA data and bulk-purified miRNA are collected from Zilionis et al. study (GSE127465) and Petriv et al.^21^ study, respectively. Single cell mRNA PDAC data was obtained from Peng et al.^22^ and lung data is obtained from Kim et al.^32^(GSE131907). mRNASeq Human Cell Landscape^7^ (HCL) is utilized for the miRSCAPE validation and the application of the global analysis.

### Model Construction and Performance Evaluation

Fig. 1A illustrates the overall miRSCAPE pipeline. We used the Extreme Gradient Boosting (xgboost) library in R language^37^ to predict the miRNA expressions using all genes’ expression values as features. We applied Grid Search for hyperparameter optimization to find the optimal parameters in the range of parameters listed in Supplementary Table S6. Given a large cohort of paired bulk mRNA and miRNA RNAseq data, we evaluated the model performance based on 5-fold cross-validation. Model accuracy was evaluated in two ways: (i) within each test sample, we quantified the Spearman correlation between the predicted and observed expression across all miRNAs. (ii) for each individual miRNA, we quantified the Spearman correlation between the observed and the predicted expression values across the test samples. For validation on single-cell data, we relied on cases where both sc-mRNA and sc-miRNA profiles are available for the same cell types. We then predicted cell type-specific miRNA profiles (using the given scRNA profiles) and then estimated the fold-difference in expression values for each miRNA across pairs of cell types; we did this using both the predicted and the observed miRNA expression values; fold-differences are estimated using the limma package in R^38^. Finally, we quantified the model accuracy as the cross-miRNA Spearman correlation between the observed and predicted fold-differences, or as Spearman correlation across miRNAs between the predicted and the observed expression within a cell type.

### Application of the model to scRNA-seq data

To apply a bulk-trained model to scRNA-seq data, we first applied the standard Seurat^34^ global-scaling normalization method. Within a single cell type, or cluster, we generated 50 bootstrapped ‘pseudo-bulk’ data by sampling without replacement 80% of the cells and pooling their expression values. This is done in order to deal with dropout noise prevalent in scRNA-seq data. To be able to apply the model trained on bulk data to the pseudo-bulk data, we ensured that the feature values are (0-1)-normalized in each sample both when training and when applying the model to pooled scRNA-seq data.

When comparing the predicted miRNAs (which have the same sample-wise expression distribution as bulk miRNA) with observed cell type miRNA, we ensured that the observed cell type data follows the same sample-wise distribution, by doing a Gaussian transformation to match the bulk miRNA distribution.

### Functional enrichment analysis

All biological functional analyses are performed with clusterProfiler^39^ package in R. Figures are generated using the ggpubr package in the R.

## Acknowledgments

This research was partially supported in part by the Intramural Research Program of the NIH, NCI, Cancer Data Science Lab. The results here are in part based upon data generated by the TCGA Research Network: https://www.cancer.gov/tcga. We thank Stefan Muljo, Shan Li, Piyush Agrawal, and Sarthak Sahoo for their valuable feedback.

## Supplementary Material

### Supplementary Results

#### 1. Comparison of performances using only the known Targets vs all genes

A primary mechanism by which a miRNA affects its target mRNA is via mRNA degradation, which would imply a negative correlation between miRNA and the mRNA. However, this expected relation may not hold given that miRNAs can also stabilize their target as well as may inhibit translation without any effect on the mRNA. Therefore, we assessed the extent to which a miRNA’s activity is informed by the expression of its known target genes relative to other genes. To this end, we first compared the fraction of all genes and the fraction of known targets that are highly correlated with a given miRNA. In three sample cohorts (CCLE, Colon cancer, Breast cancer), we observe that the miRNA exhibits a significantly higher fraction of correlation with all genes compared with the known targets suggesting that non-target genes may contribute significantly to predicting a miRNA’s activity (Supplementary Fig S1).

Next, we directly compared the cross-validation predictability of each miRNA using either only the experimentally known targets or using all genes. Again, as expected from the above, in all three cases, broadly for all miRNAs, the prediction accuracy using all genes is substantially greater than that achieved based on known targets alone (Supplementary Fig S1). We therefore decided to use all genes to predict miRNAs.

**Fig S1:**
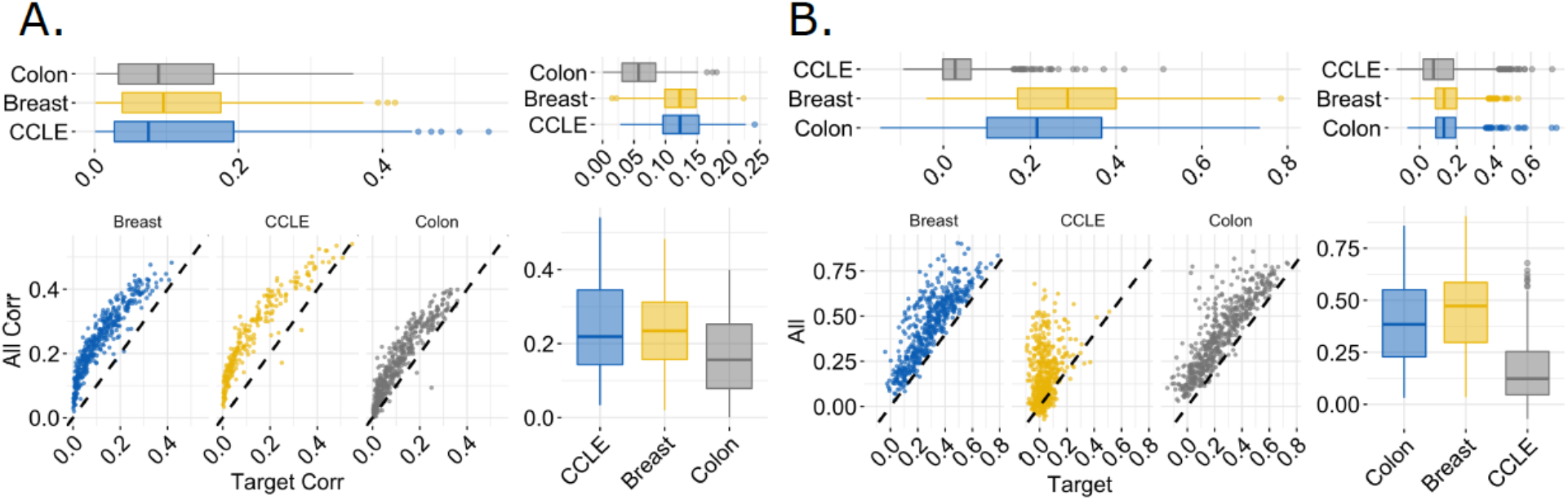
**A. Expression correlation between a miRNA and its known targets versus all genes.** The fraction of experimentally validated known targets (x-axis) and all genes (y-axis) that are correlated with a given miRNA with abs (Spearman rho) > 0.2. CCLE, TCGA colon, and TCGA breast. The top left box plot shows the distribution of the fraction of the highly correlated target genes whereas the box plot on the bottom right shows the distribution of the fraction of the highly correlated all genes. The top left box plot shows the difference between two analyses. **B. miRSCAPE performance when all genes or only experimentally known target genes are used.** Cross-validation prediction accuracy (correlation between observed and predicted expression values in test samples) for each miRNA using either all genes or only experimentally determined targets. The top left box plot shows the distribution of miRSCAPE accuracy when only experimentally known target genes are used whereas the box plot on the bottom right shows the distribution of miRSCAPE accuracy when all genes are used. The top left box plot shows the difference between two analyses. In all cases, prediction accuracy is substantially higher when using all genes as features.

#### 2. Analysis of tissue-specific gene features

Here we compared the tissue-specific models for a given miRNA that is highly predictable among 10 different tissues. For a given miRNA μ, and a pair of tissues T1 and T2 where μ is highly predictable (rho > 0.8), we identified the important features for μ, as reported by XGBoost, unique to each tissue. We then checked whether the tissue-specific important features for μ had a higher expression in the tissue where they were deemed important relative to the other tissue where they were not. Across 1,968,990 (μ,T1,T2) triplets we tested, we found that on average tissue-specific important features had 1.6-fold greater expression in the tissue where they were deemed important relative to the other tissue.

#### 3. Substantiation of miRSCAPE model using knock-out data

Loeb et al. ^40^ have reported genes that are differentially expressed (DEG) upon miR-155 knockout in CD4 T cells^41^ (GSE41241). We expect that the important features underlying the miRSCAPE model of miR-155 in hematopoietic system should either be differentially expressed upon miR-155 knockout or be correlated with the DEGs (because any machine learning model cannot distinguish between correlation and causation). We checked whether features deemed important (IFG) in the miRSCAPE model of miR-155 are correlated with the DEGs. Toward this, we tested whether IFGs exhibit a greater correlation (across our bulk training samples) with DEGs than do non-IFGs. This was indeed the case (Wilcoxon p-value < 0.05), validating the IFGs detected by miRSCAPE.

#### 4. Application of miRSCAPE to lung cancer

Focusing on the lung epithelial cells, miRSCAPE recapitulates the results of Yamada et al.^42^, showing upregulation of miR-21 in lung tumor epithelial cells. Furthermore, miR-200 family is known to inhibit tumor growth^43^ and is downregulated in the lungs of human patients and mouse model of pulmonary fibrosis linked with lung adenocarcinoma^44^. Consistently, miRSCAPE identifies miR-200 among the most down-regulated miRNAs in primary lung tumors relative to the normal samples.

Comparing metastatic and non-metastatic lung adenocarcinomas, Sun et al.^45^ found five miRNAs to be upregulated in the metastatic tumors, among which miR-210 was included in our dataset. miRSCAPE successfully identifies miR-210 among the top 20 upregulated miRNAs in the brain metastasis compared to the primary tumor in epithelial cells. Additionally, miR-145 is known to be downregulated in LUAD patients with brain metastasis^46^. Consistently, miRSCAPE identifies miR-145 to be downregulated in LUAD patients with brain metastasis, specifically in the epithelial cells.

Similar to epithelial cells, miR-210 is also upregulated in the myeloid and the lymphoid cells of the primary tumor relative to normal lung samples. However, in contrast to the epithelial cells, miRSCAPE predicts miR-21 as down-regulated in the myeloid cells of the tumor. This may represent a normal immune response consistent with a previous finding that miR-21 inhibition reduces the proportion of myeloid-derived suppressor cells in lung cancer^47^. Interestingly, miR-21 is also known to promote proliferation in AML^48^. Furthermore, miRSCAPE identified miRNA-100 among the top upregulated miRNAs in tumor myeloid cells, and like miR-21, miR-100 is a known oncomir for AML^49^, suggesting a link between miRNA functions in the lymphoid cells across contexts.

MicroRNA miR-328 represents another interesting case. It promotes myeloid differentiation^50^. Eiring et al.^51^ have shown loss of miR-328 in CML. We observed the downregulation of miR-328 in the lung tumor myeloid cells. However, in contrast, miR-328 is upregulated in the lymphoid cells of the brain metastatic tumors, consistent with Arora et al.^52^ again, suggesting a complex, context-specific role of miRNAs revealed by miRSCAPE.

Analyzing the peripheral blood lymphocytes of pulmonary sarcoidosis (linked with lung cancer), Kiszałkiewicz et al.^53^ found significant upregulation of miR-222 and significant downregulation of let-7f in patients compared to controls. miRSCAPE recapitulates these findings in lung cancer lymphoid cells vs normal lung lymphoid cells.

#### 5. Application of miRSCAPE to Human Cell Atlas

Thum et al. list 52 upregulated and 40 downregulated miRNAs between human fetal and adult heart tissue. Among those 46 overlapped genes, miRSCAPE recapitulates 26 upregulated and 10 downregulated miRNAs corresponding to a concordance rate of 78.26%. (Fig 5D). Tang et al.^54^ investigate miRNAs in matched human fetal and adult organs (heart, kidney, liver, lung). They report that miR-21 is overexpressed in the fetal lung. Consistently, miRSCAPE predicts an increase for miR-21 in the lung. MiRSCAPE estimates the expression for miR-26a is upregulated in fetal lung and kidney compared to adult. Tang et al. also observe an overexpression for miR-26a in the fetal lung and kidney. Additionally, they suggested that miR-125b is functional both at the fetal and the adult stages of kidneys. Consistently, we also observe a high level in miR-125b in both fetal and adult stages of kidney with no difference in expression.

Comparing cardiac remodeling in fetal and adult rats, Yan et al.^55^ detected miR-199a and miR-21 to be upregulated in adults. This is consistent with our findings. Moreover, Zhang et al.^56^ identified 18 and 7 miRNAs that respectively increase and decrease with age in mice. Of these miRSCAPE consistently identified 14 and 4 miRNAs that respectively increase or decrease in adult cardiac cells relative to fetal cardiac cells.

## Supplementary Figures

**Fig S2:**
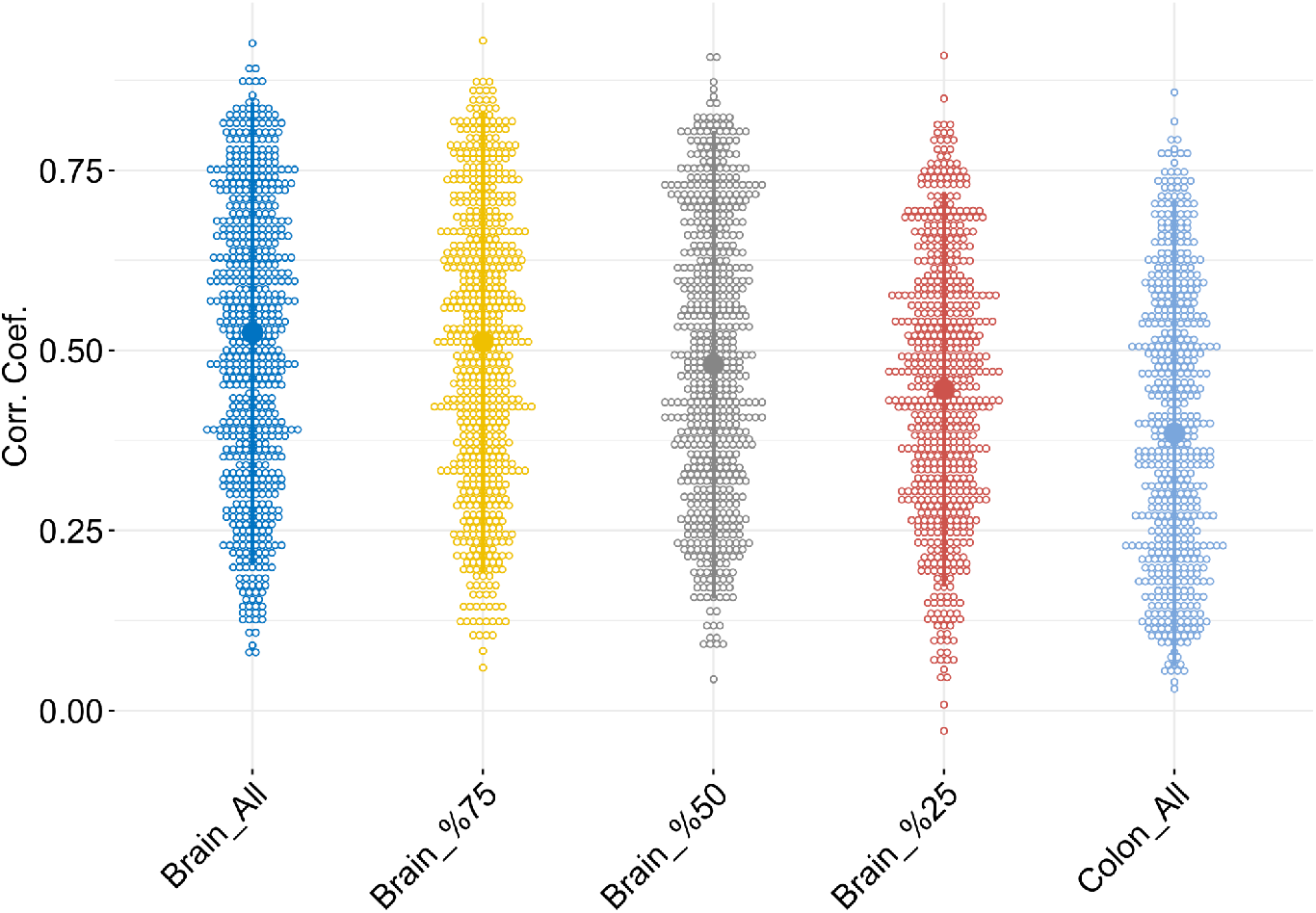
Effect of the sample sizes on the miRSCAPE prediction. Performance of the miRSCAPE is provided for all (n = 507), 75% (n = 380), 50% (n = 253), 25% (n = 126) of the brain samples and all colon (n = 443) samples are used. miRSCAPE performance is slightly affected by the sample size since a mild accuracy decrease when part of the samples is utilized in brain cohorts. However, the sample size is not the only performance factor as the lowest accuracy is obtained on the colon (n = 443) samples.

**Fig 3:**
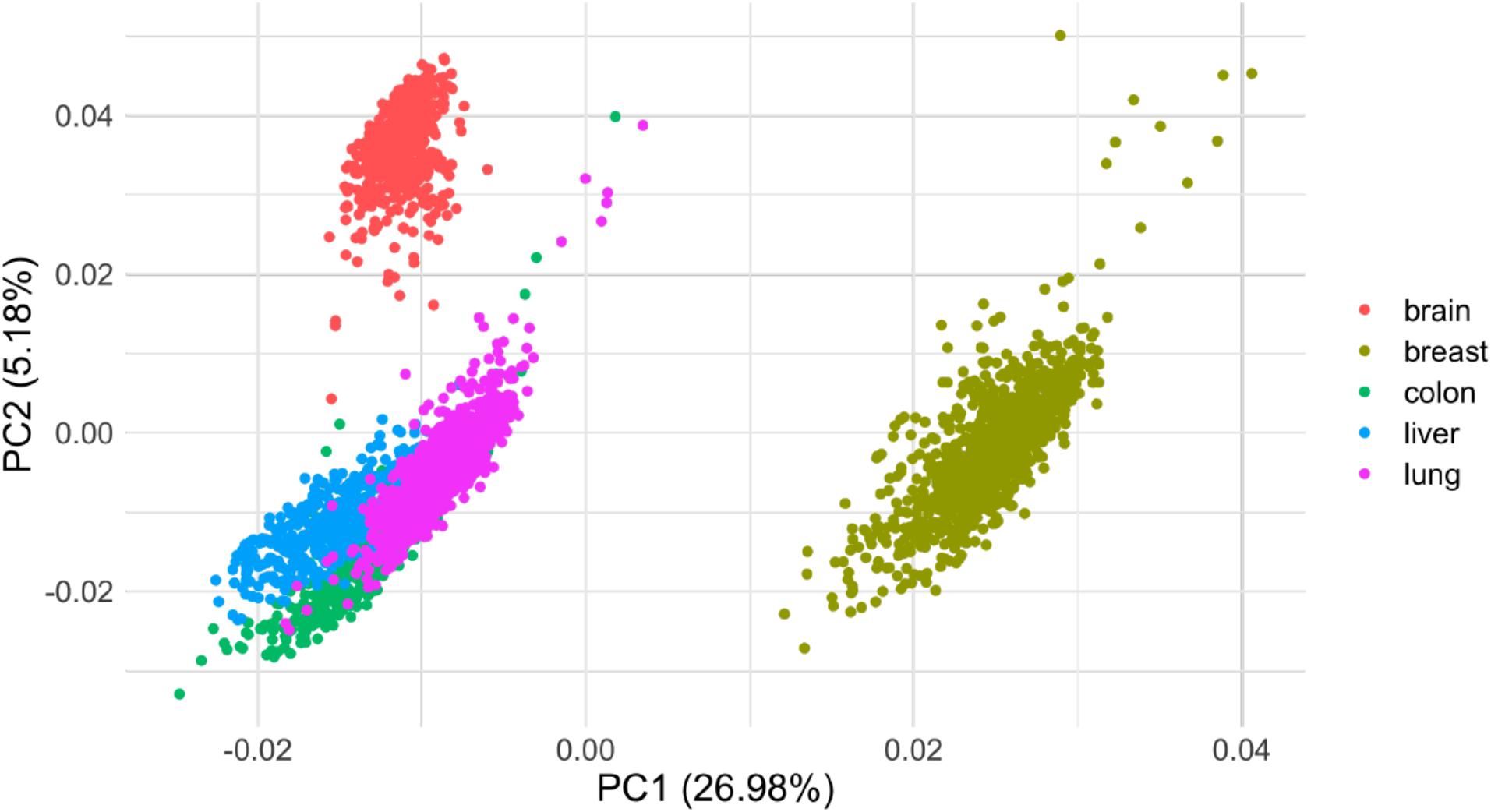
Principal component analysis (PCA) plot for the mRNA expression profiles of 5 cancer types.

**Fig S4:**
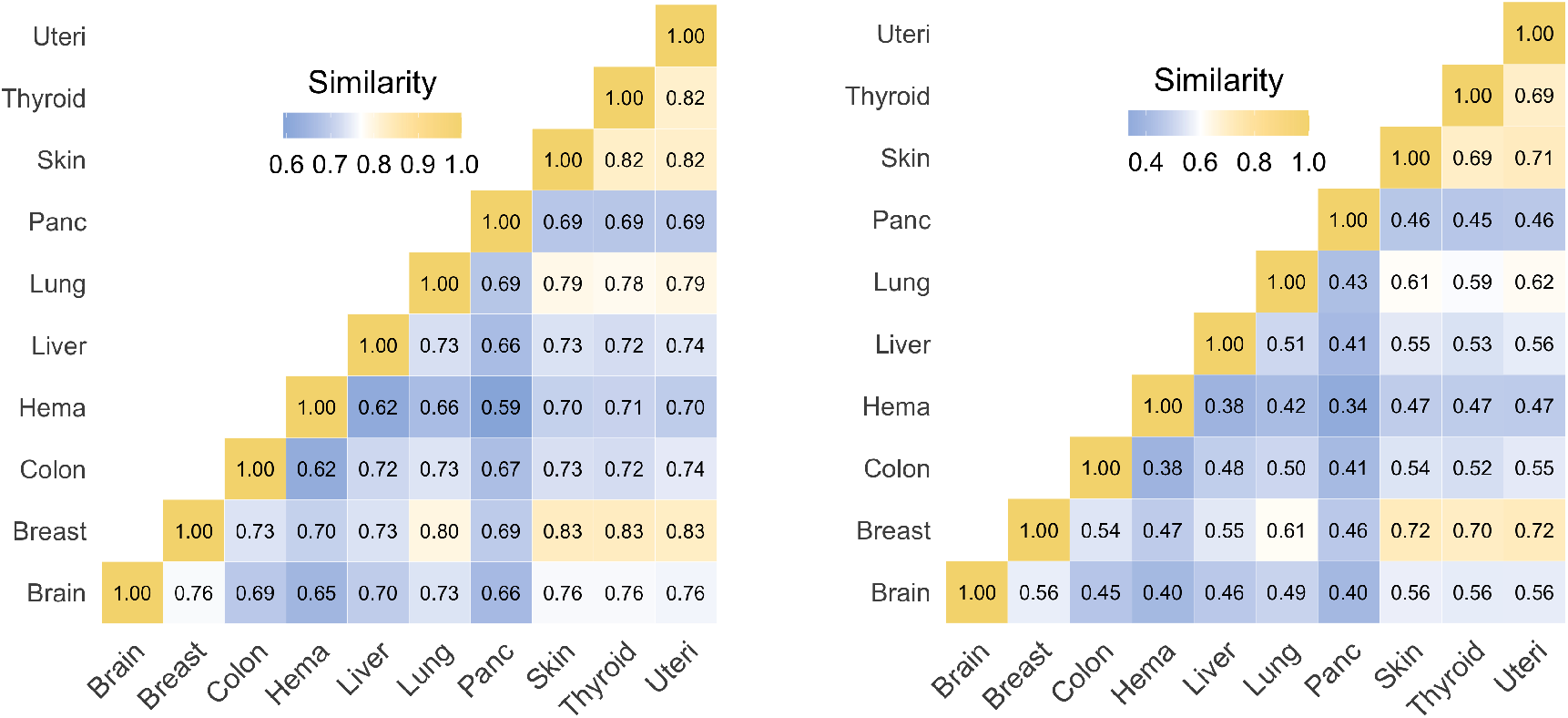
Cross-tissue similarity (Jaccard index) in important features - those deemed important for at least 50% (left) and 80% (right) of the miRNAs in each tissue.

**Fig S5:**
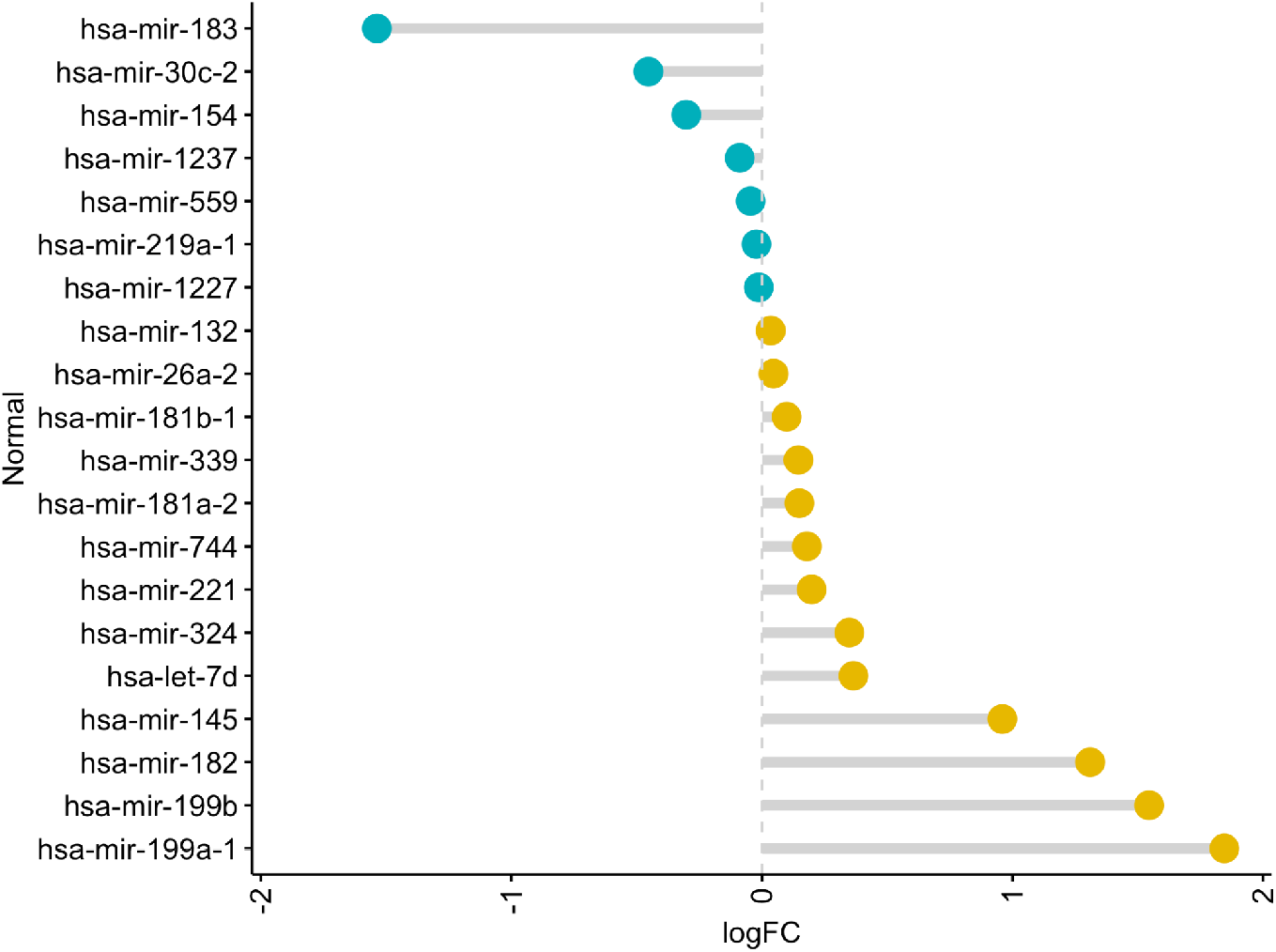
Fold change (x-axis) distribution of the predicted miRNAs (y-axis) differentially expressed in PDAC malignant cells versus pooled acinar and ductal type 1 cells. miRNAs whose fold change is consistent with Mazza et al. are shown.

## Supplementary Tables

**Table S1:** Number of samples and number of miRNAs covered in this analysis for the cancer and normal data

**Table S2:** Top 20 upregulated and downregulated miRNAs’ logFC and p-values in the malignant cells relative to acinar and ductal Type I cells individually as well as relative to pooled acinar and ductal cells in PDAC.

**Table S3:** The top 20 upregulated and downregulated miRNAs’ logFC and p-values relative to primary tumors versus normal samples and also the metastatic tumors versus the primary tumors separately in epithelial, lymphoid, and myeloid cells. T: Primary Tumor; N: Normal; M: Metastatic tumor.

**Table S4:** The top 20 upregulated and downregulated miRNAs’ logFC and p-values for each of the 56 different cell types relative to all other cell types.

**Table S5:** The top 20 upregulated and downregulated miRNAs’ logFC for the matching tissues between the fetal and the adult tissues.

**Table S6:** The utilized list of parameters and their ranges in this analysis.

## Notes

### Competing Interest Statement

The authors have declared no competing interest.

